# Environmental Uncertainty Affects Movement and Space-use in Sheep

**DOI:** 10.1101/2023.08.10.552758

**Authors:** Sarah T. Bartsch, William H. E. J. van Wettere, Simon C. Griffith, Stephan T. Leu

**Affiliations:** The University of Adelaide, School of Animal and Veterinary Science, Australia; The University of Adelaide, School of Animal and Veterinary Science, Davies Livestock Research Centre, Australia; Macquarie University, School of Natural Sciences, Australia; University of New South Wales, School of Biological, Earth and Environmental Sciences, Australia

**Keywords:** Foraging, information, movement ecology, space-use, uncertainty, ungulates

## Abstract

Animals constantly experience periods of uncertainty due to seasonal changes in food distribution. The changing climate results in more variable weather patterns, which in turn alter environmental conditions, and can result in resource distribution being less predictable in space and time. How animals respond to these uncertain conditions, in particular the changing distribution of food resources, remains largely unclear and is an important question in the field of movement and animal ecology. Here we used an experimental approach to study how Merino sheep (*Ovis aries*) responded to different levels of environmental uncertainty in a drought-impacted region of the Australian arid zone. Sheep were unfamiliar with the experimental resource distribution at the start and progressively decreased their uncertainty (i.e., increased their environmental knowledge) when discovering an increasing number of foraging patches. We tracked 50 sheep with GPS collars (1 location every 15 sec) and deduced their movement and space use behaviour. When environmental uncertainty decreased, individuals moved more directionally (greater step length, smaller turn angles) and moved greater distances per day. They also had larger daily home ranges but rested in similar areas on consecutive nights (similar displacement, with the exception when five patches were discovered). Our study demonstrates how an arid zone, free-ranging ungulate adjusts its movement and space use behaviour as it gains environmental information in order to forage efficiently during periods of uncertainty. Our study provides important insights into how animals cope with variable environments and different levels of uncertainty.

## INTRODUCTION

Most animal species move through space to locate food. However, an animal’s movements can be constrained or altered by movement barriers, such as habitat fragmentation or uncertainty about where to find resources (Armansin et al., 2020; Nathan et al., 2008). The way animals respond behaviourally to uncertainty caused by a changing environment can determine how effectively the animal can locate crucial resources (Cunningham et al., 2015; Dall et al., 2005; Fagan et al., 2013; Herzing et al., 2017). For example, elephants respond to drought by travelling further distances for food which results in higher calf survival rates and ultimately improves fitness (Foley et al., 2008). Drought and increased temperatures, which can be exacerbated by climate change (Trenberth, 2011) can cause differences to the distribution and availability of food resources over time and space (Frank and McNaughton, 1992). The often unpredictable transformations in habitat require the animal to gather new information about the resource distribution. These habitat changes can induce exploratory movement and animals have to travel further to locate resources (Kefi et al., 2007; Papageorgiou et al., 2021). The increased energetic costs and difficulty of doing so, could lead to mortality and ultimately local species extinctions (Foley et al., 2008; Paniw et al., 2019; Trisos et al., 2020; Urban, 2015).

Animals acquire environmental information and locate resources using different sensory cues and spatial memories (Edwards et al., 1996; Lee et al., 2006). When an animal has limited knowledge about its environment, i.e., when uncertainty is high, the animal can make less informed movement decisions, which can result in energetically costly movement patterns. In the context of foraging, this may ultimately lead to non-optimal foraging strategies with a reduced likelihood of encountering the best available resources. Individuals show different movement patterns when they are initially unaware of the location of their goal compared to when the location is known and re-visited (Krakauer and Rodŕíguez-Gironés, 1995; Rook et al., 2005). For example, a slow local search, characterised by short step lengths (low speeds) and many turns, covers a small area in high detail. It is used to gain information about the local environment prior to leaving an area. Once the decision has been made to leave, such as when resource availability becomes low, the step length is longer, and the movement is more directional. This covers a larger area at lower detail in order to find food that is further away (Bartumeus et al., 2005; Moorter et al., 2013; Venter et al., 2017). However, it can be more difficult to perceive resources than when using a local search.

The typical space an animal uses, i.e., its home range, contains critical resources for survival and reproduction, such as food, shelter and mates (Senft et al., 1987). If the resource needs of an individual increase, the home range size can increase (Foley et al., 2008; Larter and Gates, 1990; Papageorgiou et al., 2021; Ullmann et al., 2018). For example, vulturine guineafowl increase their ranging distances and home range size after periods of drought to meet individual resource needs and support their population (Papageorgiou et al., 2021). If movement patterns change in order to find more dispersed resources, it is conceivable that this can also affect home range size.

In addition to environmental conditions, inter-or intra-species competition can influence changes in movement behaviour (Barta and Giraldeau, 1998). For example, when competition for food is high, individuals often move to alternative, or more distant food sources to mitigate the direct costs of competition. Displacement through competition is more likely when food sources are rare and can be monopolised (Herfindal et al., 2019; Wignall et al., 2020). When uncertainty increases, for instance when animals are exploring a new habitat, the number of food patches that have been discovered is low. Hence, during periods of uncertainty, food sources may be perceived as rare, resulting in high levels of competition irrespective of the true abundance of the resource.

Using an experimental approach, we investigated how animals modify their movement and space use behaviour as information about the environment (food locations) is acquired. Few studies have been able to experimentally test hypotheses on the effect of uncertainty in mammals at a defined temporal and spatial scale. In our study we used sheep, because of the amenability with which they can be exposed to experimental treatments, across large spatial scales. Sheep can be enclosed in defined areas thereby controlling the environment each individual experiences. Furthermore, sheep can be easily captured, and sex and age composition can be controlled.

We predicted that when animals experience higher levels of uncertainty, they travel (i) faster and (ii) longer distances to find additional food patches and to reduce conspecific competition at the patches already found. Furthermore, we predicted that (iii) the daily displacement (distance between the first and last location of the day) increases. If the animals travel further to find food, it is conceivable that they do not return to the same locations at night where they started the day, and instead use different resting areas. As a consequence of the larger travel distance, we further predicted that (iv) home ranges are larger during periods of uncertainty, and that (v) movement paths would be more tortuous (larger turn angles) when searching for food patches.

Using sheep as a study species in controlled environmental experiments allowed us to experimentally investigate movement and space-use in response to uncertainty at a large and realistic scale, thereby also providing important comparative insight for other mammals, including wildlife species. Movement and space use measures provide insight into how animals respond to changing and uncertain food resource distribution (Bartumeus et al., 2016). It is crucial to understand how individuals and ultimately, species can cope with the changing environments, in particular since human activity is exacerbating environmental change (Gross, 2018).

## METHODS

### Experimental resource distribution and level of uncertainty

We conducted the study at Fowler’s Gap Arid Zone Research Station, New South Wales, Australia (31°05′S, 141°43′E). The study area is characterised as arid chenopod shrubland, with predominantly blue bush (*Maireana* spp.). The general area and our study paddock (six x one kilometre with a water trough in one corner) were affected by severe drought over consecutive years. This resulted in limited grass available to forage, and the experimentally provisioned food patches represented a patchy and high-value resource.

We selected eight food locations within the study area that met the following criteria: (i) more than 200 metres away from the fence line, (ii) at least 1500 metres from the water trough (located in the Northwest corner of the paddock), and (iii) at least 400 metres apart from other food locations. At the start of the experiment, we placed two bales of lucerne hay (approximately weighing 20 kilograms each) at each of those eight locations. Then we released the sheep into the paddock near the water trough. We placed two extra bales of lucerne at the water trough to assist the sheep with acclimatisation after being handled.

We conducted the experiment over 15 days. We excluded three days of data from the analysis which were collected during days of replenishment or relocation of food patches, this left 12 days of data for analysis (Table 1). Food patches were replenished after every three days. After six days, eight new food patches were established following the same criteria as above. All new locations were at least 400m apart from any previous location. We replenished or relocated patches whilst the animals were sighted at the water trough and not within view of the food patches.

**Table 1.**
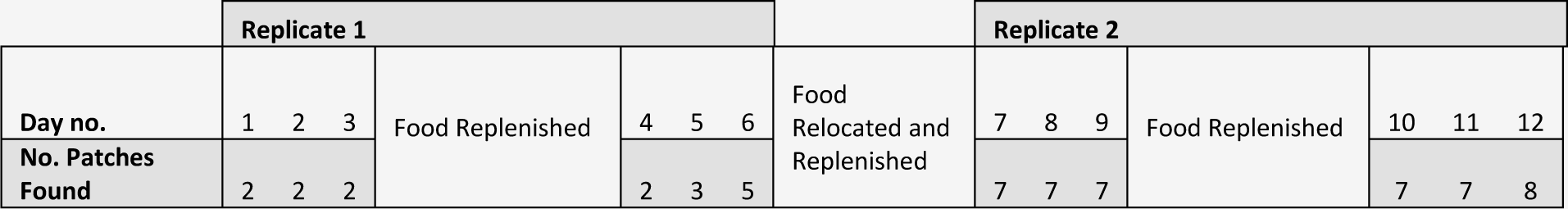
Cumulative number of food patches discovered throughout the study, out of eight possible patch locations.

### Tracking animals with GPS collars

We recorded the movement and space use behaviour of 50, two-year-old non-pregnant female sheep (ewes). Each individual was fitted with a global positioning system (GPS) collar weighing 700g (MobileAction GPS, i-gotU GT-120, and Core Electronics battery CE04381). The GPS collars recorded the location of each sheep every 15 seconds for 16 hours during daylight hours between 0430 hours and 2030 hours local time. The collars were synchronised to enable all sheep locations to be recorded simultaneously.

### Ethical Note

At the start of the study all sheep were moved into a yard and each individual was fitted with a GPS collar. The collar represented 1.9 percent of the mean body mass of our study sheep (mean = 37.3 kg, SE = 0.6 kg), and 2.7 percent of the lightest animal. Earlier studies have shown that collars do not affect natural behaviour (Hulbert et al., 1998), and we did not notice any unusual behaviours after collaring and prior to data collection. When we removed all GPS collars at the end of the project, we did not notice any skin irritation, and the sheep were released back into the paddock. All sheep were treated using procedures formally approved by the University of New South Wales Animal Care and Ethics Committee and Macquarie University Animal Ethics Committee in compliance with the Australian Code of Practice for the Use of Animals for Scientific Purposes (approval numbers 18/19B and 3249, respectively).

### GPS data processing

The location data were processed in R (v4.1.2; R Development Core Team, 2021), using the methods described in Leu et al. (2021). In short, we removed GPS locations if they (i) were recorded with less than three satellites, (ii) visually fell outside the known paddock boundaries, or (iii) exceeded the animal’s maximum movement speed, or (iv) deviated from the movement path (Bjørneraas et al., 2010), indicated by high movement speeds and large turn angles (above 120 degrees). We used the function *vmask* from the R package *argosfilter* for the movement speed filter which uses two previous and two following locations. We used a speed threshold of 5.34 kilometres per hour, which represents the movement speed of a sheep during a simulated predator attack (Manning et al., 2014). The 15-second recording interval can vary slightly among records and GPS units as units sometimes take longer to record a location. We accounted for this, by interpolating all GPS locations of all sheep such that they perfectly fitted 15-second intervals. We used the function *na.approx* from the R package *zoo*, to interpolate the data. The interpolation process assumes straight and constant movement between locations. The locations were usually only offset by a few seconds. Hence, we expect that the assumption of straight movement within those seconds is reasonably well met. The function can also estimate missing locations using the known locations before and after the gap. However, we only did this if a maximum of two consecutive locations were missing, i.e., a maximum gap of 30 seconds, to avoid estimated locations that deviated too greatly from the empirical path (Leu et al., 2021).

### Determining the level of environmental uncertainty

We considered a patch to be discovered if a sheep had been within a 20-metre distance of the food location. We used the *distGeo* function to calculate this. We used the number of discovered food patches as a proxy for the level of uncertainty. The level of uncertainty for each day was determined by the maximal number of food patches that were discovered until the end of this day. With each additional patch being discovered, sheep had greater environmental knowledge. We initially thought that relocating all food patches would start a second period of uncertainty. However, the sheep were fast to discover the new food patches, and the post-relocation period in fact reflected greater resource knowledge (Table 1).

### The effect of uncertainty on movement and space use behaviour

For each individual, we calculated the step length as the distance between successive GPS locations, (metres travelled per 15 seconds). We also calculated the total distance travelled per day (in kilometres) by summing up the step lengths of that day. Next, we determined the net displacement of each individual as the distance between the first and last location of the day. Turn angles were calculated as a measure of bearing and travel changes along a path. Home range was determined as 100% minimum convex polygon. We used the R function *prepData* of the package *moveHMM*, *distGeo* of the package *geosphere*, and *mcp* of the package *AdehabitatHR* for our calculations. Separate linear mixed models were used to determine the relationship between the number of discovered patches and our movement and space use measures. The mean step length, total distance travelled, displacement, home range and turn angles were the dependent variable in separate models, and the number of patches discovered (eight levels), and the period (two levels, before and after the resource relocation) were fixed factors. Sheep ID was included as a random variable to account for inter-individual differences. An example of the mixed model structure is: mean step length ∼ no. patches discovered + period + (1|ID)

We used the *lmerTest* R package for our statistical analysis, which uses Satterthwaite’s method to deduce P-values for the fixed effects as well as degrees of freedom. The *Performance* R package was used for visual inspections of residual plots to determine that the data fulfilled the assumptions of homoscedasticity and normality. An alpha level of 0.05 was selected to identify significant results.

## RESULTS

### The effect of uncertainty on movement and space use behaviour

The dataset consisted of a total of 2,916,901 GPS locations after processing. After running our separate linear mixed models, and because the number of discovered patches was a fixed factor, we used post hoc tests to determine whether our measures of movement behaviour and space use changed significantly with additional patches discovered (Note, patch 4 and 5 were discovered on the same day, patch 6 and 7 were also discovered on the same day). The step length significantly increased with each one to two additional patches discovered (Fig 1A, Table 2). The total distances travelled per day also significantly increased as patches were discovered (Fig 1B, Table 3). The displacement distance was not affected by the number of patches discovered, and the animals rested in similar areas on consecutive nights (Fig 1C, Table 4). The only exception and significant difference in displacement was when five patches were discovered, and animals rested approx. 680 metres further apart than when three patches were found. Home range size significantly increased as the number of discovered patches increased (Fig 1D, Table 5). Finally, the turn angle decreased with the number of patches discovered, that is, movement became more directional with increased knowledge. This was not the case at the beginning of the experiment, and turn angles were not significantly different when two and three patches were discovered (Fig 1E, Table 6.).

**Fig. 1.**
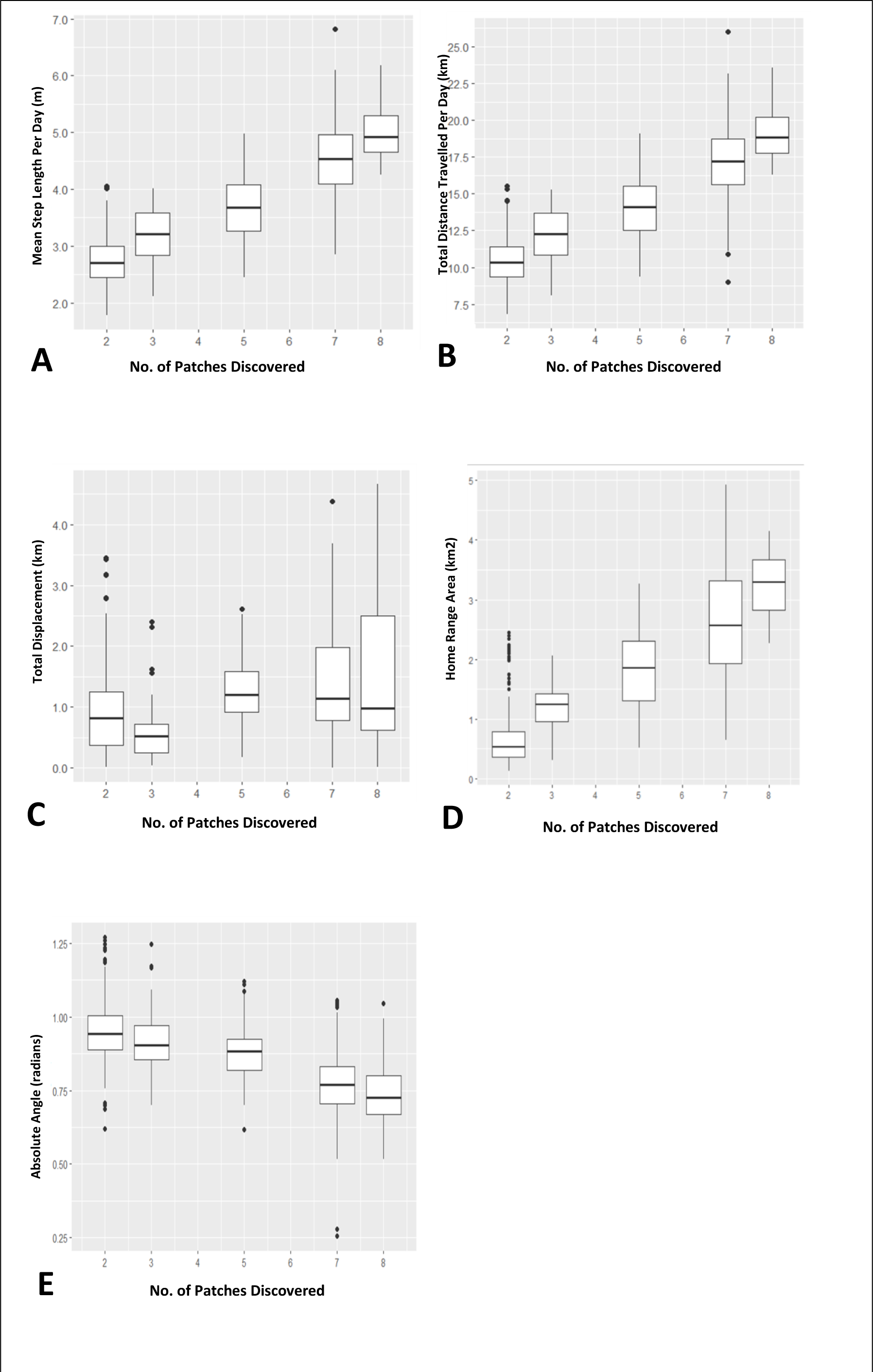
Sheep movement characteristics differ in response to the number of food patches discovered, a proxy for uncertainty. More patches discovered indicate lower levels of uncertainty of environmental conditions. The experimental setup included eight separate resource locations. The panels show that with a *greater* number of patches discovered **A** the mean step length increased (step length = metres moved per 15 seconds), **B** total daily distances travelled were higher, **C** daily displacement increased when 5 patches were discovered compared to 3 patches, **D** home ranges were larger, **E** mean absolute turn angles were smaller. The data for patch 4 and 6 are not shown separately because it was discovered during the same day as patch 5 and 7, respectively.

**Table 2.** Mean step length in relation to uncertainty, i.e., the number of food patches discovered.

| Increasing no. patches discovered | Estimate (m) | SE | DF | *t*-Value | *p*-Value |
| --- | --- | --- | --- | --- | --- |
| 2 to 3 | 0.438 | 0.078 | 546.000 | 5.611 | **<0.001** |
| 3 to 5 | 0.502 | 0.099 | 546.000 | 5.087 | **<0.001** |
| 5 to 7 | 0.843 | 0.076 | 546.000 | 11.026 | **<0.001** |
| 7 to 8 | 0.478 | 0.076 | 546.000 | 6.254 | **<0.001** |

Significant effects are shown in bold.

**Table 3.** Distance travelled per day in relation to uncertainty, i.e., the number of food patches discovered.

| Increasing<br>no. patches<br>discovered | Estimate<br>(km) | SE | DF | *t*-Value | *p*-Value |
| --- | --- | --- | --- | --- | --- |
| 2 to 3 | 1.675 | 0.295 | 546.000 | 5.686 | **<0.001** |
| 3 to 5 | 1.905 | 0.373 | 546.000 | 5.111 | **<0.001** |
| 5 to 7 | 3.044 | 0.289 | 546.000 | 10.545 | **<0.001** |
| 7 to 8 | 1.975 | 0.289 | 546.000 | 6.841 | **<0.001** |

Significant effects are shown in bold.

**Table 4.** Displacement per day in relation to uncertainty, i.e., the number of food patches discovered.

| Increasing<br>no. patches<br>discovered | Effect (km) | SE | DF | *t*-Value | *p*-Value |
| --- | --- | --- | --- | --- | --- |
| 2 to 3 | -0.341 | 0.126 | 546.000 | -2.705 | 0.054 |
| 3 to 5 | 0.679 | 0.160 | 546.000 | 4.255 | **<0.001** |
| 5 to 7 | 0.113 | 0.124 | 546.000 | 0.912 | 0.892 |
| 7 to 8 | 0.088 | 0.124 | 546.000 | 0.713 | 0.953 |

Significant effects are shown in bold.

**Table 5.** Home range per day in relation to uncertainty, i.e., the number of food patches discovered.

| Increasing<br>no. patches<br>discovered | Effect<br>(km<sup>2</sup>) | SE | DF | *t*-Value | *p*-Value |
| --- | --- | --- | --- | --- | --- |
| 2 to 3 | 0.510 | 0.106 | 546.000 | 4.834 | **<0.001** |
| 3 to 5 | 0.592 | 0.133 | 546.000 | 4.437 | **<0.001** |
| 5 to 7 | 0.820 | 0.103 | 546.000 | 7.931 | **<0.001** |
| 7 to 8 | 0.653 | 0.103 | 546.000 | 6.320 | **<0.001** |

Significant effects are shown in bold.

**Table 6.** Mean absolute turn angles per day in relation to uncertainty, i.e., the number of food patches discovered.

| Increasing<br>no. patches<br>discovered | Effect | SE | DF | *t*-Value | *p*-Value |
| --- | --- | --- | --- | --- | --- |
| 2 to 3 | -0.027 | 0.011 | 546.000 | -2.475 | 0.098 |
| 3 to 5 | -0.043 | 0.014 | 546.000 | -3.092 | **0.018** |
| 5 to 7 | -0.112 | 0.011 | 546.000 | -10.294 | **<0.001** |
| 7 to 8 | -0.035 | 0.011 | 546.000 | -3.221 | **0.012** |

Significant effects are shown in bold.

## DISCUSSION

We found clear evidence for an effect of uncertainty on movement and space use behaviour. Here, we used the number of discovered experimental food patches as a measure of knowledge gained, i.e., the more patches that were discovered, the lower the level of uncertainty. Movement was faster and more directional as uncertainty decreased. Animals also moved longer distances per day and had larger home ranges. With environmental knowledge, displacement distance only significantly increased once, on the day when 5 patches had been discovered compared to 3 patches. At the end of the day, returning to an area close to where they started their day irrespective of the level of uncertainty regarding food patches, would suggest that sheep repeatedly use similar rest areas, independent of their foraging movement.

The changes in movement behaviour in relation to the level of uncertainty is supported by the work of Bartumeus et al. (2016), suggesting that the level of environmental uncertainty relates to different search strategies which are associated with particular movement characteristics. For instance, initial spatial explorative behaviours resulting from a state of uncertainty could be described as investigative movements used by the animal to then select the most efficient movement style (Bartumeus et al., 2016; Bracis et al., 2015). Gathering information or exploring the environment is a process of learning which results in changes to behaviour and foraging decisions as the individual gains experience (Krakauer and Rodríguez-Gironés, 1995).

As predicted, sheep moved with larger turn angles when uncertainty was high, and more directionally as knowledge of patch locations increased. Severe reorientation is often the result of randomised movement when faced with extreme uncertainty within a locality (Bartumeus et al., 2005). These substantial directional changes can decrease as environmental experience is acquired and new movement behaviours are adopted to create a more informed, systematic, and efficient area scan. Reduced turn angles are also consistent with moving between and revisiting known patches. However, contrary to our predictions, sheep travelled faster, covered longer distances, and had larger home ranges on a given day when environmental knowledge increased. A potential explanation for this finding is that after some patches were discovered and sheep learned that high-quality food patches existed in the landscape, they continued to sample the environment to gain information. Hence, they did not remain in the vicinity of resource patches that they had already discovered but moved around to find more patches (Bartumeus et al., 2016; Dumont and Petit, 1998). Furthermore, the movement characteristics may also suggest that the sheep continued to move back and forth between known food locations when more and more patches were discovered (Bracis et al., 2015). Hence, animals may leave a discovered food patch before it was depleted and return at a later time (Seidel and Boyce, 2015). Consistent with this notion, sheep may have used an area-restricted search strategy when uncertainty was high. This search strategy is effective when resource locations are wide, scarce, and patchy, or when animals lack pre-existing knowledge of resource targets outside of their explored area (Dorfman et al., 2022). When using an area-restricted search strategy, animals scan a local area, using slow, tortuous, short movements to exploit a resource patch before leaving. Once a few patches were discovered, the sheep quickly adjusted their movement. Similarly, Hewitson et al. (2005) suggested that sheep adjust their foraging behaviour in response to the predictability of the food item location within a patch. When food item locations within a patch were variable and hence less predictable, sheep increased their environmental sampling within the patch before leaving it (Hewitson et al., 2005).

Exploring a new habitat and moving larger distances is linked with an increasing home range size. For example, home range size has been demonstrated to increase in elephants and vulturine guineafowl when resources are limited (Foley et al., 2008; Papageorgiou et al., 2021). This would suggest that home range size increased with a lack of knowledge of food locations, and when fewer food patches had been discovered. In contrast, we found that sheep had small daily home ranges when uncertainty was high, but home ranges increased in size with the number of discovered food patches. As before, and consistent with the movement measures, this could be explained by a continuation of exploring the area, or by sheep moving between the increased number of already discovered patches.

At the start of the experiment, sheep were not familiar with the resource distribution and their search strategy likely initially reflected a landscape without high-quality patches. The new resource environment required the animals to change their strategy to locate the food patches. Halfway through the experiment, we relocated all food patches, with the intention of increasing the level of uncertainty again. However, the sheep very quickly found the new patches, which may be due to the animals already having adopted a search strategy and movement style that was optimal for finding the dispersed food (Ullmann et al., 2018). The food relocation reflected the same resource distribution although in different locations and would not require a change of search strategy (Dorfman et al., 2022).

## Conclusion

In this study, we showed how different levels of uncertainty affect the movement and space use behaviour of free-ranging sheep. Using an experimental approach, we were able to empirically quantify the level of environmental uncertainty that the sheep experienced, which is usually difficult to achieve in free-living animals. Here, we assumed that on a given day the level of uncertainty was similar for all individuals, which is a useful simplification. Further investigations that integrate individual levels of uncertainty and group composition would provide very nuanced knowledge of foraging strategies in heterogeneous groups. Nevertheless, our study provides deep insight into how a free-ranging ungulate modulates its movement and space use, which is indicative of its foraging behaviour as knowledge about its environment increases. Wild animals constantly experience periods of uncertainty, due to seasonal changes in food distribution but also due to climate change, which can result in dynamic and unpredictable changes in resource availability and distribution in space and time. Using sheep to obtain knowledge of the behavioural responses to these ecological challenges is an important step toward understanding the impact that environmental change could have on wild, free-ranging mammals.

## ACKNOWLEDGEMENTS

We thank Keith Leggett (Fowler’s Gap research director) for project support. We also thank Garry Dowling (Fowler’s Gap station manager), Mark Tilley, and Derek Kennedy for their support during sheep handling and mustering. This work and STL were supported by an Australian Research Council DECRA Fellowship [DE170101132]. The Queensland Tertiary Admissions Centre RRESP Scholarship and the University of Adelaide for the School of Animal and Veterinary Sciences Honours Scholarship supported STB.

## AUTHOR CONTRIBUTIONS

**Sarah T. Bartsch:** Conceptualization, Formal analysis, Data curation, Writing original draft, Funding acquisition **William H. E. J. van Wettere:** Writing – Review and Editing, Supervision **Simon C. Griffith:** Conceptualization, Writing – Review and Editing **Stephan T. Leu:** Conceptualization, Data curation, Writing – Review and Editing, Supervision, Funding acquisition

## DECLARATION OF INTEREST

We declare no conflicts of interest.

## Notes

### Competing Interest Statement

The authors have declared no competing interest.

